# Association between land use and composition of amphibian species in temperate Brazilian forest remnants

**DOI:** 10.1101/2021.02.17.431642

**Authors:** Roseli Coelho dos Santos, Diego Anderson Dalmolin, Diego Brum, Mauricio Roberto Veronez, Elaine Maria Lucas, Alexandro Marques Tozetti

## Abstract

We evaluated the influence of landscape configuration on the diversity of anurans in Atlantic Forest remnants in southern Brazil. As natural habits provide better conditions for the survival of amphibians, we expected to find more diverse communities in areas with more forest cover. We sampled tadpoles in 28 breeding sites distributed in seven forest remnants. We recorded 22 anuran species and richness varied from 6 to 12 species between sites. Most of the recorded species were not forest specialists, except for *Boana curupi* and *Crossodactylus schmidti*. There was a significant overlap in the species composition between all remnants, and the Generalized Linear Mixed Model indicated that landscape use did not affect species richness. The PERMANOVA showed that forest and livestock farming explained the dissimilarity in the composition of the communities. One possible explanation for this is that the remnants are surrounded by a relatively well-preserved landscape, which offers favorable conditions for the maintenance of local populations and homogenizes species composition across the sampling sites. The lack of any strong association between tadpole species richness and land use suggests that anurans are primally affected by habitat characteristics that are detected only on a fine-scale analysis.

## Introduction

The increase of the human population generates demands such as larger areas for agriculture and livestock farming, as well as more roads and buildings, which leads to increased consumption of natural resources. Consequently, humans promote a variety of landscape modifications (Gururaja et al. 2008; Zhou et al. 2017). When natural habitat is replaced by roads, buildings or other urban facilities, many ecological interactions can be affected, such as predation, competition and host/parasite relationships (Nomura et al. 2011; Laufer et al. 2015; Preuss et al. 2020; Santos et al. 2020). These changes can vary according to the intensity in which habitat is lost and air, soil and water are polluted (Riley et al. 2005; Brand et al. 2010).

One of the most studied negative effects of agriculture on wildlife is its role as a source of contamination by pesticides and herbicides (Koumaris and Fahrig 2016). However, when a forest is replaced by agriculture or cattle pasture, besides all issues listed below, there will be also a sensible reduction in both habitat complexity and landscape heterogeneity (Machado et al. 2012; Saccol et al. 2017), which favors a gradual reduction in species diversity (McKinney 2008; Barrows and Allen 2010). As the human occupation of a landscape increases, the proximity between natural habitats and urban areas reduces, causing a series of indirect (and less studied) impacts on the fauna, such as light pollution (Dias et al. 2019), noise pollution (Pellet et al. 2004) and road kills (Diniz and Brito 2015). In sum, the facts listed above highlight the relevance of constant monitoring changes in landscape induced by human occupation.

Landscape evaluation is a powerful tool to access the risks of biodiversity deployment and, consequently, supports environmental management. At the species level, landscape configuration affects fauna displacement (Fahrig and Nuttle 2005), which is a baseline for testing models of metapopulation and metacommunities. In general, species with low displacement ability, such as amphibians, are more sensitive to landscape changes (Schmutzer et al. 2008; Dixo and Metzger 2010; Diniz and Brito 2013; Cayuela et al. 2015). Most amphibians breed in aquatic sites, and changes in landscapes can restrain access to ponds or streams (Becker et al. 2007, 2010; Machado et al. 2012; Cayuela et al. 2015; Saccol et al. 2017; Dalmolin et al. 2019). This can promote morphological and physiological changes in tadpoles and adults (Costa et al. 2017) and reduce reproductive potential, causing changes in the communities’ structure and composition (Berriozabal-Islas et al. 2018; Dalmolin et al. 2020), as well as population declines or local extinctions (Marsh and Trenham 2001; Rothermel 2004; Goutte et al. 2013). Therefore, amphibians are considered good models for ecological studies on understanding the impact of landscape changes on wildlife. The monitoring of amphibian populations has been filling the gaps in our understanding of how landscape changes affect species diversity (Becker et al. 2007; Pillsbury and Miller 2008; Nomura et al. 2011; Collins and Fahrig 2017) and the physiology and morphology of amphibians (Costa et al. 2017; Berriozabal-Islas et al. 2018). This highlights the importance of studies that include a greater variety of altered or natural habitats and conduct measurements in the surroundings of the sampled habitats.

The Brazilian Atlantic Forest is highly fragmented, with more than 97% of its fragments being smaller than 250 ha (Ribeiro et al. 2009; Zanella et al. 2012). The Atlantic Forest is a biodiversity hotspot (Myers 2000; Mittermeier et al. 2005) and harbors a high anuran richness and endemism (Haddad et al. 2013). In the southern Atlantic Forest, most of the original forest areas were replaced by agriculture, pasture, silviculture and urban areas (Ribeiro et al. 2011). Studies on amphibians in this region have associated species diversity with characteristics of habitat heterogeneity (Gonçalves et al. 2015; Knauth et al. 2018; Figueiredo et al. 2019), but few have analyzed wider scales that considered the effects of the landscape on species composition. Landscape analyses allow the detection of changes regarding land use and occupation in scales that are compatible with the species or groups in question (Hamer and Parris 2011), helping define local and regional conservation strategies.

Many studies have focused on the role of size and connectivity of forest fragments on species diversity. However, less attention is given to the role of surrounding environments generated by human occupation. In this study, we evaluated the influence of the surrounding landscape configuration on the diversity of anurans in forest remnants in southern Brazil. Our analysis included quantification of different types of land use surrounding these forest remnants. As natural habitats provide better conditions for the survival, reproduction and maintenance of amphibian populations (Collins and Fahrig 2017), we expected to find more diverse communities in areas with more forest cover.

## Material and methods

### Study site

We conducted this study in Atlantic Forest remnants in southern Brazil (Fig. 1; Table S1). Forest remnants consist of mixed ombrophilous forest and seasonal forest with a variety of surrounding matrix including livestock pastures, agriculture, silviculture and urban areas (SOS Mata Atlântica and INPE 2008; Ribeiro et al. 2009; Pillar and Vélez 2010). The climate is subtropical, with annual temperatures varying from 16 ºC to 24 ºC and rainfall varies from 1600 to 2200 mm annually (Alvares et al. 2013). We selected seven remnants based on the following criteria: a) presence of well-preserved forest (public or private protected areas); b) similar climatic conditions; c) altitude between 300 m and 900 m above the sea level; d) similar topography. We sampled seven remnants distributed across 10.500.000 ha, between the coordinates 22º30’ to 33º45’ S (latitude) and 48º02’ to 57º40’ W (longitude).

**Fig. 1.**
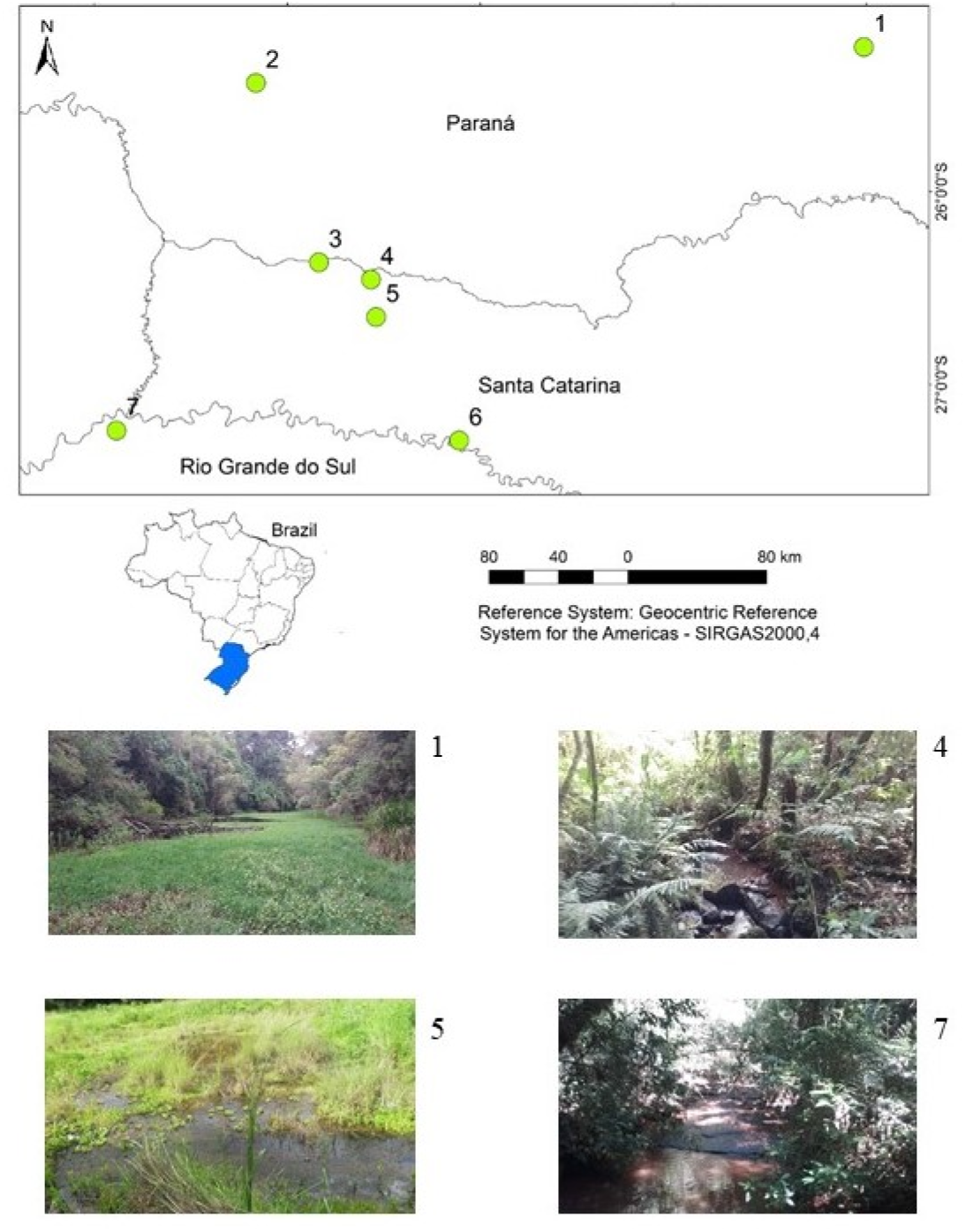
Forest remnants and sampling sites of amphibian collection conducted between October 2018 and March 2019 in southern Brazil. 1 = Parque Estadual de Vila Velha, Ponta Grossa/PR; 2 = Parque Estadual Rio Guarani, Três Barras/PR; 3 = Reserva Privada Enele, São Lourenço do Oeste/SC; 4 = Parque Estadual das Araucárias, São Domingos/SC; 5 = Reserva Privada Quebra-Queixo, São Domingos/SC; 6 = Parque Estadual Fritz Plaumann, Concórdia/SC; 7 = Parque Estadual do Turvo, Derrubadas/RS. The detail depicts sampling points of the sampling units – 1 = Pond in Parque Estadual de Vila Velha; 4 = Stream in Parque Estadual das Araucárias; 5 = Pond in Reserva Privada Quebra-Queixo; 7 = Stream in Parque Estadual do Turvo

In each remnant, we performed tadpole sampling in water bodies. We sampled a total of 28 water bodies used as breeding sites (13 ponds and 15 streams). We sampled all possible waterbodies including permanent and temporary ponds and streams to access a greater variety of habitats and their tadpole communities (Melo et al. 2017). Pond size varied from 71.20 to 2087 m^2^ (787.27 ± 561.33). In streams, we sampled 100-m-long sections with a width varying from 0.60 to 3.50 m (1.76 ± 0.79). The depth of the ponds varied from 0.30 to 1.50 m (0.8 ± 0.40) and of streams from 0.15 to 0.70 m (0.45 ± 0.12; Appendix, Table S1). We performed a time-limited sampling effort.

### Tadpole sampling

Tadpole sampling occurred from October 2018 to March 2019, which corresponds to the anuran breeding season and is and the most favorable period for detection of tadpoles (Both et al. 2008; Santos et al. 2008). Tadpole sampling was performed from 8:00 AM to 6:00 PM using a 3-mm^2^-mesh dip-net (Heyer 1976). Sweepings were systematized and consisted of scouring the margins of the pond (Vasconcelos and Rossa-Feres 2005; Santos et al. 2007; Both et al. 2009; Bolzan et al. 2016). In the streams, some samples were also made using small dip nets to sample narrow spaces between rocks (adapted from Jordani et al. 2017). Sampling was performed once in each pond or stream. Each sample consisted of one hour of sweeping efforts through the greatest possible variety of microhabitats for one hour.

Immediately after capture, tadpoles were euthanized by immersion in a solution of 2% lidocaine, following Brazilian Regulations (CONCEA 2018), and subsequently transferred to absolute ethanol. In the laboratory, tadpole species were determined with the aid of a stereomicroscope and identification keys (Machado and Maltchik 2007; Gonçalves 2014). The collection and manipulation of specimens were authorized by ICMBio (#64962), state organs (IAP/PR #33/2018, IMA/SC #01/2019, SEMA/RS #37/2018) and the Animal Ethics Committee of Universidade do Vale do Rio dos Sinos (CEUA-UNISINOS # PPECEUA08.2018).

### Landscape assessment

We assessed land use based on the analysis of satellite images (Landsat 8 multispectral images, sensor Operational Land Imager from United States Geological Survey). We classified the images from their vectorization in the software ArcGIS version 10.3, considering a 250-m-radius buffer for each sampled water body. Buffer size was defined based on the estimative of the size of habitats used by anurans (Semlitsch and Bodie 2003; Smith and Green 2005; Tozetti and Toledo 2005; Canessa and Parris 2013). All images were captured in the current sampled year (2019) and presented the minimal cloud cover available and without significant radiometric noise. We performed the following stages of image preprocessing: 1) geometric corrections due to the inherent geometric distortions in images collected in distinct moments, by georeferencing these images; 2) atmospheric corrections aiming to reduce the interference of atmospheric scattering on the images (Soares et al. 2015); and 3) mosaicing and highlight of the different images in each moment aiming to reduce seasonal effects on the image’s visual aspect. Preprocessing was conducted with the software ENVI, version 5.51. After the pre-processing stages, we defined the categories of land use and occupation with adjustments based on field observations. We were able to identify the following categories (landscape variables) of land use: (1) Agriculture (cultivated areas, with soybean, corn or wheat); (2) Aquatic environments (streams, artificial and natural ponds); (3) Forest (native forest formations); (4) Livestock pastures (extensive livestock farming); (5) Urban area (buildings). The polygons regarding each cover type were projected for the reference system SIRGAS 2000, Universal Transverse Mercator (UTM) projection, zone 22S, and areas were calculated in km^2^.

### Data analysis

The comparison of species richness was based on rarefaction curves (interpolation and extrapolation method) representing standardized measures of individual abundance (Chao and Jost 2012). Confidence intervals (95%) associated with the curves were calculated using the Bootstrap method (50 randomizations). These analyses were performed using the iNEXT program (iNterpolation and EXtrapolation) (Chao and Hsieh 2016).

We performed a Principal Components Analysis (PCA) to check which variables were more associated with the landscape configuration of each remnant (and its community). The PCA was performed using a water bodies matrix and a landscape matrix with a 250 m buffer.

We tested the influence of landscape variables (Appendix, Table S2) on species richness by using generalized linear mixed-effects models (GLMMs). We included water bodies as a random variable. We evaluated the significance of each explanatory variable by using the “ANOVA” function. The full model was analyzed using the “glm” function of the lme4 package (Bates et al. 2015), in R v.3.6.0 (R Core Team 2019).

We assessed the differences in species composition between remnants (β-diversity) and the relationship with landscape variables by using Permutational Multivariate Analysis of Variance (PERMANOVA) with 999 permutations (Borcard et al. 2011; Magurran and McGill 2011). Additionally, we used a Non-Metric Multidimensional Scaling (NMDS) to visualize and interpret these differences.

PERMANOVA and NMDS were performed, respectively, by using the “adonis” and “metaMDS” functions of the vegan package in R v.3.6.0 (R Core Team 2019). For analysis using landscape variables on species diversity, only 23 water bodies were used to avoid overlapping buffers used to assess land use.

## Results

We recorded 22 anuran species belonging to eight families: Bufonidae (2 species; 9.1% of the total), Hylidae (12; 54.5%), Hylodidae (1; 4.5%), Leptodactylidae (3; 13.6%), Microhylidae (1; 4.5%), Odontophrynidae (1; 4.5%), Phyllomedusidae (1; 4.5%) and Ranidae (1; 4.5%), the latter represented by the exotic species *Lithobates catesbeianus* (Table 1). Generalist species were predominant in terms of habitat use, with exception of *Boana curupi* and *Crossodactylus schimidti*, which are forest-related species. Species composition was dominated by habitat generalists (*Boana faber, Dendropsophus minutus, Lithobates catesbeianus, Physalaemus* cf. *carrizorum, P. cuvieri, Scinax fuscovarius*). The richness among forest remnants ranged from 6 to 12 species (8,72 ±2,06), and abundance from 213 to 680 individuals (408.7 ±101.6; Table 1). Interpolation and extrapolation curves evidenced that species richness differed between remnants (Fig. 2).

**Table 1.**
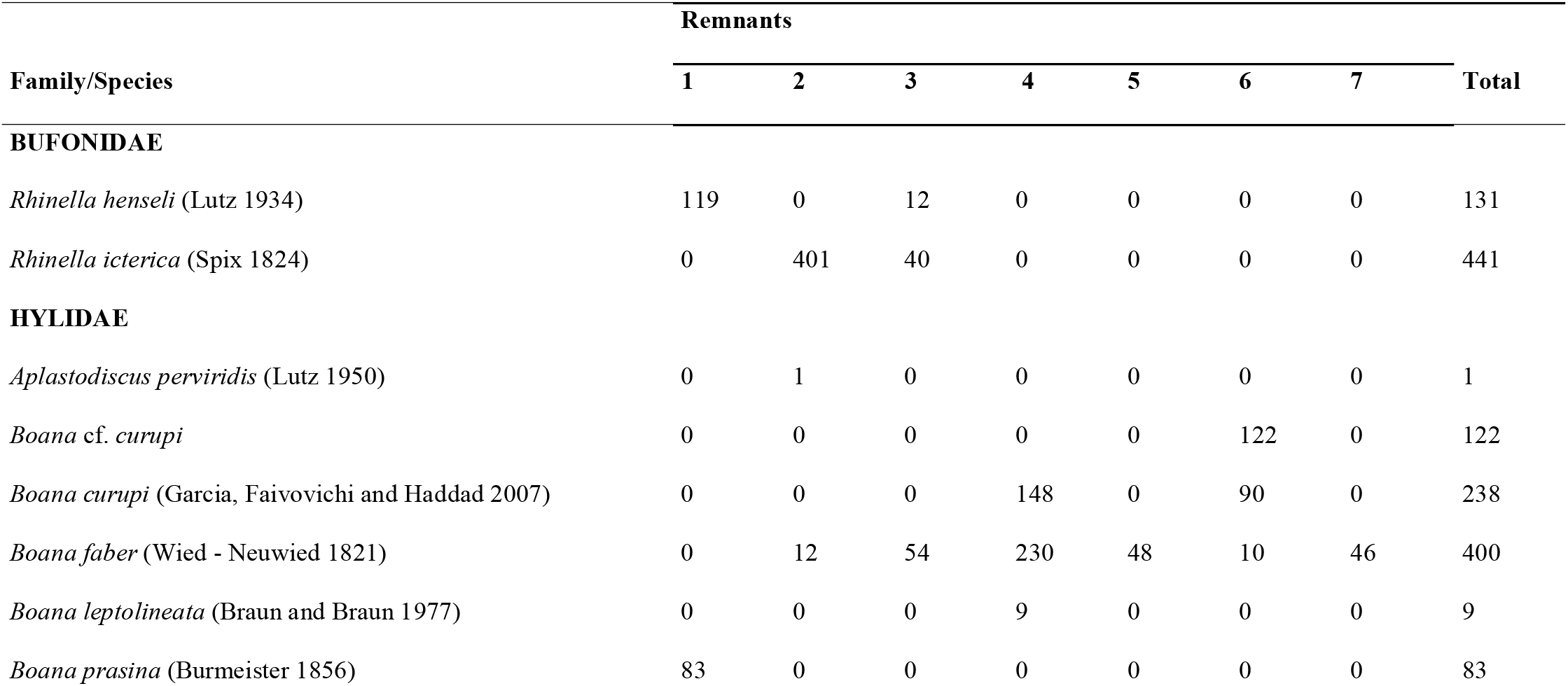
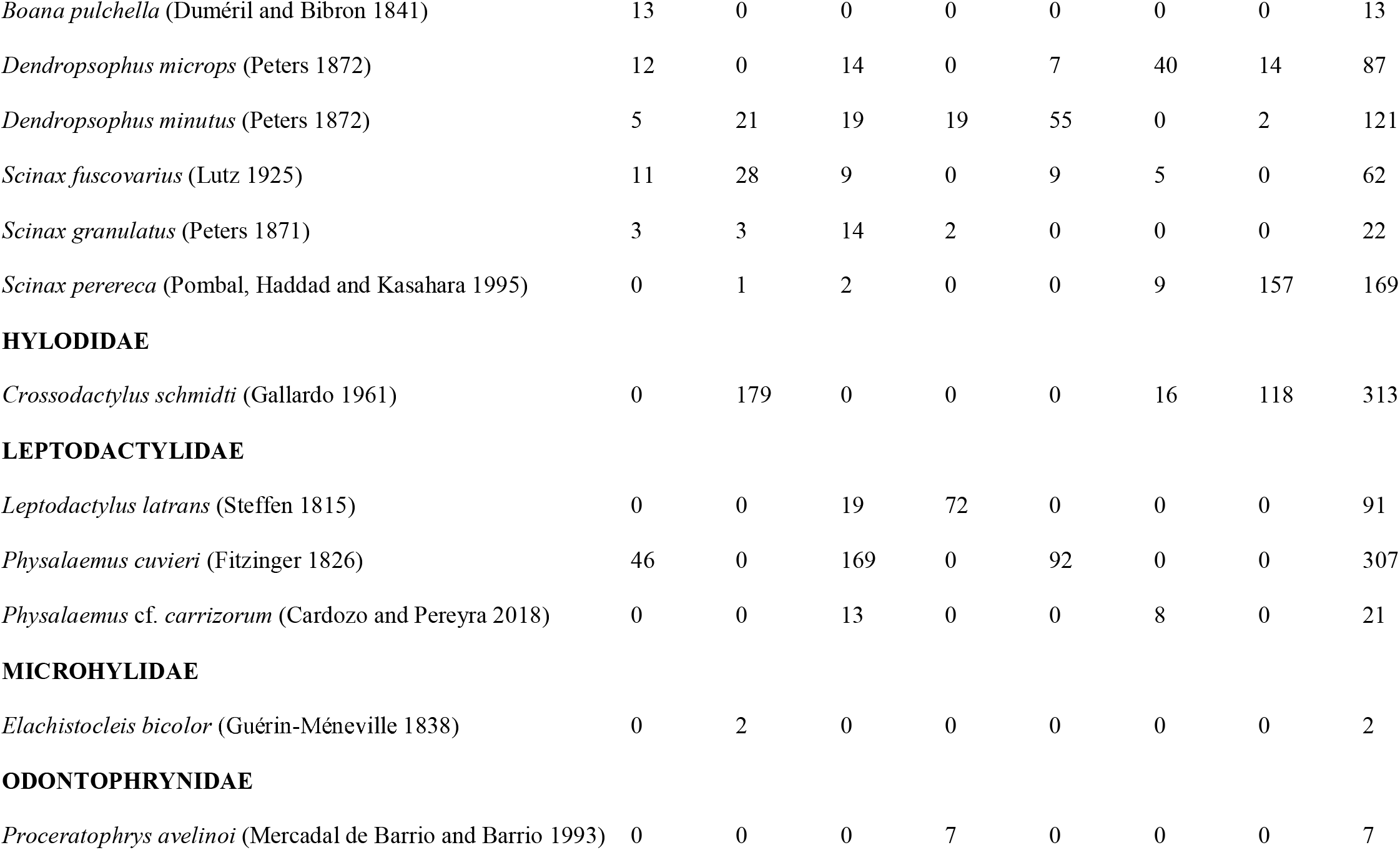
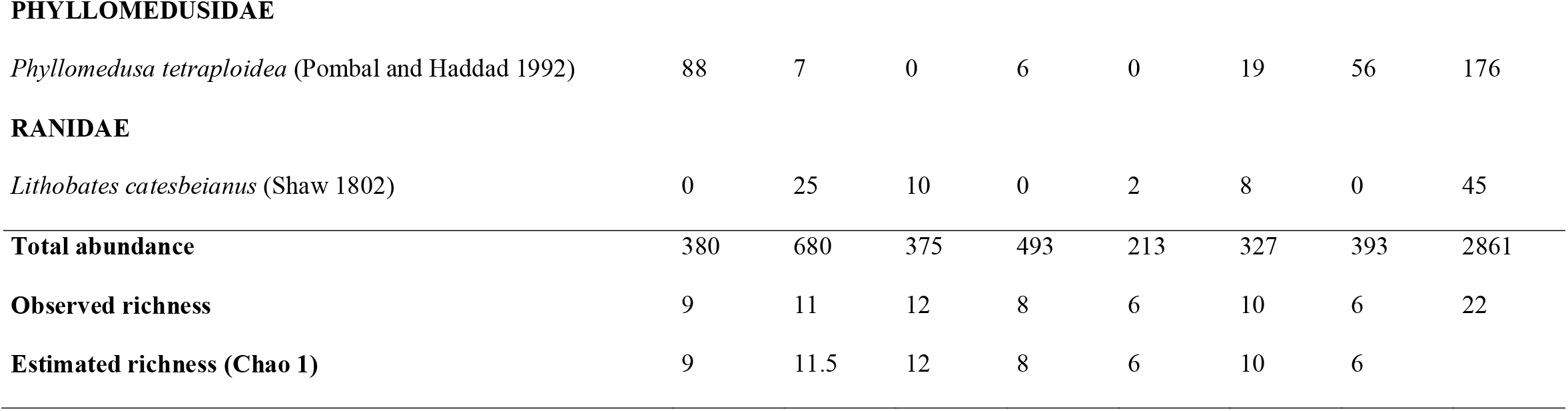
Tadpole abundance and observed and estimated richness (Chao 1) for the seven sampled remnants, in southern Brazil. Sites: 1 = Parque Estadual de Vila Velha; 2 = Parque Estadual Rio Guarani; 3 = Reserva Privada Enele; 4 = Parque Estadual das Araucárias; 5 = Reserva Privada Quebra Queixo; 6 = Parque Estadual Fritz Plaumann; 7 = Parque Estadual do Turvo

| Family/Species | Remnants |  |  |  |  |  |  | Total |
| --- | --- | --- | --- | --- | --- | --- | --- | --- |
|  | 1 | 2 | 3 | 4 | 5 | 6 | 7 |  |
| BUFONIDAE |  |  |  |  |  |  |  |  |
| *Rhinella henseli* (Lutz 1934) | 119 | 0 | 12 | 0 | 0 | 0 | 0 | 131 |
| *Rhinella icterica* (Spix 1824) | 0 | 401 | 40 | 0 | 0 | 0 | 0 | 441 |
| HYLIDAE |  |  |  |  |  |  |  |  |
| *Aplastodiscus perviridis* (Lutz 1950) | 0 | 1 | 0 | 0 | 0 | 0 | 0 | 1 |
| *Boana* cf. *curupi* | 0 | 0 | 0 | 0 | 0 | 122 | 0 | 122 |
| *Boana curupi* (Garcia, Faivovich and Haddad 2007) | 0 | 0 | 0 | 148 | 0 | 90 | 0 | 238 |
| *Boana faber* (Wied - Neuwied 1821) | 0 | 12 | 54 | 230 | 48 | 10 | 46 | 400 |
| *Boana leptolineata* (Braun and Braun 1977) | 0 | 0 | 0 | 9 | 0 | 0 | 0 | 9 |
| *Boana prasina* (Burmeister 1856) | 83 | 0 | 0 | 0 | 0 | 0 | 0 | 83 |
| *Boana pulchella* (Duméril and Bibron 1841) | 13 | 0 | 0 | 0 | 0 | 0 | 0 | 13 |
| *Dendropsophus microps* (Peters 1872) | 12 | 0 | 14 | 0 | 7 | 40 | 14 | 87 |
| *Dendropsophus minutus* (Peters 1872) | 5 | 21 | 19 | 19 | 55 | 0 | 2 | 121 |
| *Scinax fuscovarius* (Lutz 1925) | 11 | 28 | 9 | 0 | 9 | 5 | 0 | 62 |
| *Scinax granulatus* (Peters 1871) | 3 | 3 | 14 | 2 | 0 | 0 | 0 | 22 |
| *Scinax perereca* (Pombal, Haddad and Kasahara 1995) | 0 | 1 | 2 | 0 | 0 | 9 | 157 | 169 |
| **HYLODIDAE** |  |  |  |  |  |  |  |  |
| *Crossodactylus schmidtii* (Gallardo 1961) | 0 | 179 | 0 | 0 | 0 | 16 | 118 | 313 |
| **LEPTODACTYLIDAE** |  |  |  |  |  |  |  |  |
| *Leptodactylus latrans* (Steffen 1815) | 0 | 0 | 19 | 72 | 0 | 0 | 0 | 91 |
| *Physalaemus cuvieri* (Fitzinger 1826) | 46 | 0 | 169 | 0 | 92 | 0 | 0 | 307 |
| *Physalaemus* cf. *carrizorum* (Cardozo and Pereyra 2018) | 0 | 0 | 13 | 0 | 0 | 8 | 0 | 21 |
| **MICROHYLIDAE** |  |  |  |  |  |  |  |  |
| *Elachistocleis bicolor* (Guérin-Méneville 1838) | 0 | 2 | 0 | 0 | 0 | 0 | 0 | 2 |
| **ODONTOPHRYNIDAE** |  |  |  |  |  |  |  |  |
| *Proceratophrys avelinoi* (Mercadal de Barrio and Barrio 1993) | 0 | 0 | 0 | 7 | 0 | 0 | 0 | 7 |
| **PHYLLOMEDUSIDAE** |  |  |  |  |  |  |  |  |
| *Phyllomedusa tetraploidea* (Pombal and Haddad 1992) | 88 | 7 | 0 | 6 | 0 | 19 | 56 | 176 |
| **RANIDAE** |  |  |  |  |  |  |  |  |
| *Lithobates catesbeianus* (Shaw 1802) | 0 | 25 | 10 | 0 | 2 | 8 | 0 | 45 |
| **Total abundance** | 380 | 680 | 375 | 493 | 213 | 327 | 393 | 2861 |
| **Observed richness** | 9 | 11 | 12 | 8 | 6 | 10 | 6 | 22 |
| **Estimated richness (Chao 1)** | 9 | 11.5 | 12 | 8 | 6 | 10 | 6 |  |

**Fig. 2.**
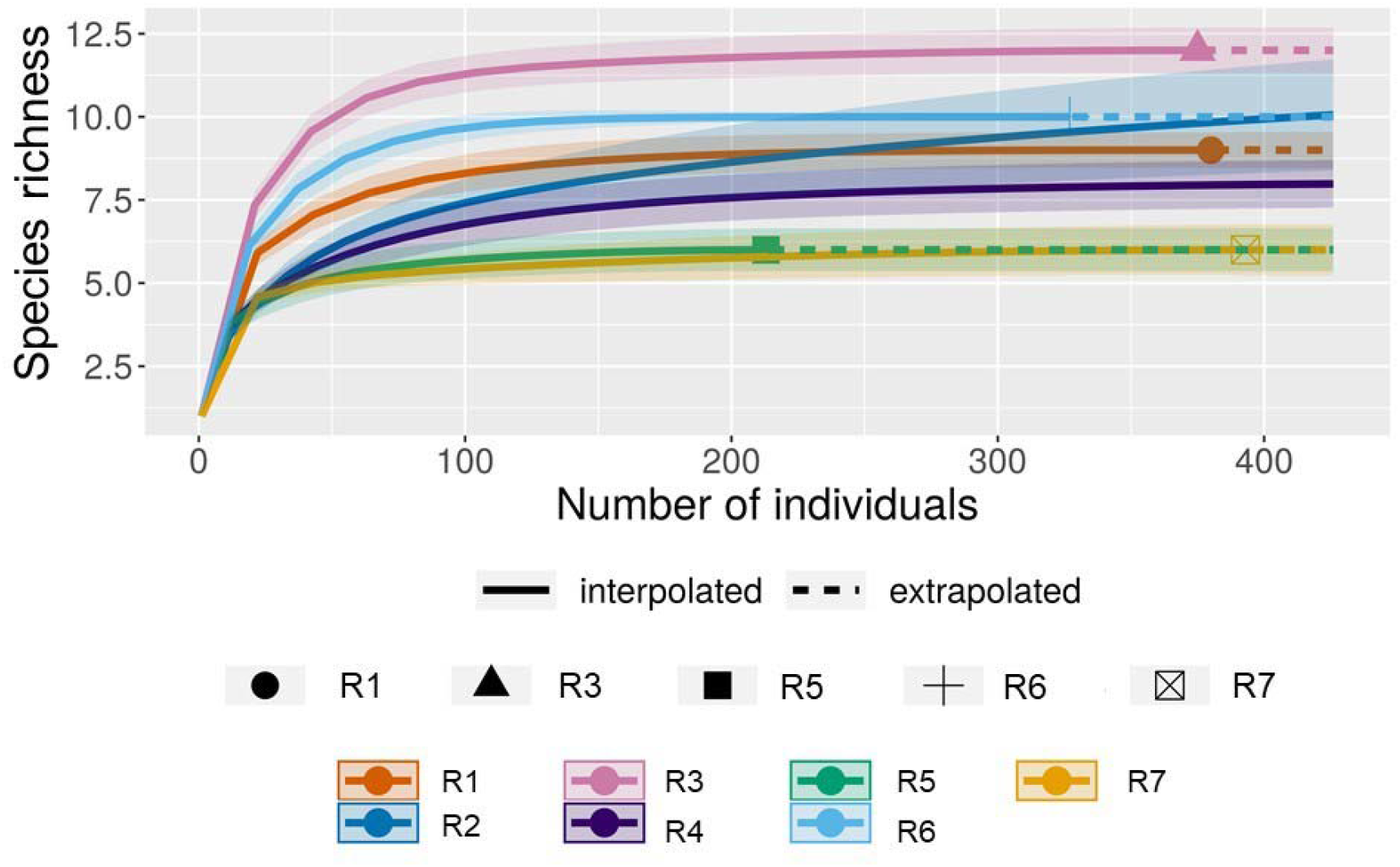
Comparison of the richness of anuran species in seven Atlantic Forest remnants in southern Brazil through rarefaction (interpolation/extrapolation) based on the number of individuals. R1 (orange) = Parque Estadual de Vila Velha; R2 (navy blue) = Parque Estadual Rio Guarani; R3 (lilac) = Reserva Privada Enele; R4 (purple) = Parque Estadual das Araucárias; R5 (green) = Reserva Privada Quebra-Queixo; R6 (sky blue) = Parque Estadual Fritz Plaumann; R7 (yellow) = Parque Estadual do Turvo

Each of the two of PCA’s axes was related to different types of land use and together, explained 62% of the variation of land use surrounding the remnants (Table S3). The presence of forest, livestock pastures and aquatic environment were more associated with axis 1, while agriculture and urbanization were more associated with axis 2. Tadpole communities presented visible segregation in the biplot, with the remnants 2, 3, 5 and 7 mainly distributed across axis 1 and remnants 1, 4 and 6, across axis 2 (Fig. 3; Table S4). However, the GLMM model showed that landscape use did not affect species richness (Table 2), with the largest amount of richness (75%) explained by a random effect (R^2^m = 0.14; R^2^c = 0.89). With exception of remnants 1 and 6, all forest remnants presented high overlap in their species composition (Fig. 4).

**Fig. 3.**
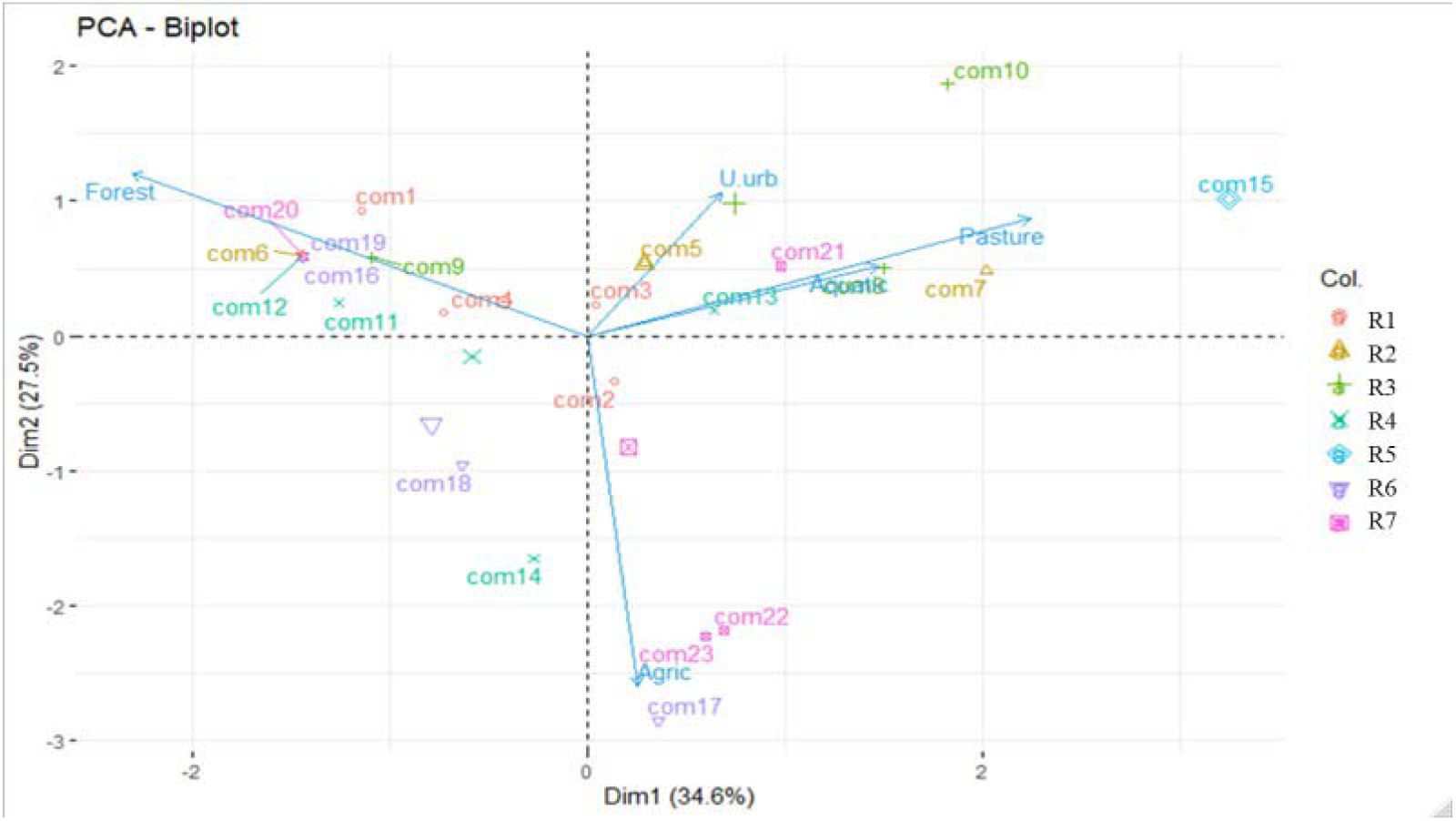
Ordination biplot with the two first axes of the Principal Components Analysis considering the association between 23 sampled breeding sites in seven remnants and the landscape variables in southern Brazil. Symbols (○, ∆, +, ×, ◊, ▼, □) represent the remnants. ○ R1 = Parque Estadual de Vila Velha; ∆ R2 = Parque Estadual Rio Guarani; + R3 = Reserva Privada Enele; × R4 = Parque Estadual das Araucárias; ◊ R5 = Reserva Privada Quebra-Queixo; ▼ R6 = Parque Estadual Fritz Plaumann; □ R7 = Parque Estadual do Turvo

**Table 2.**
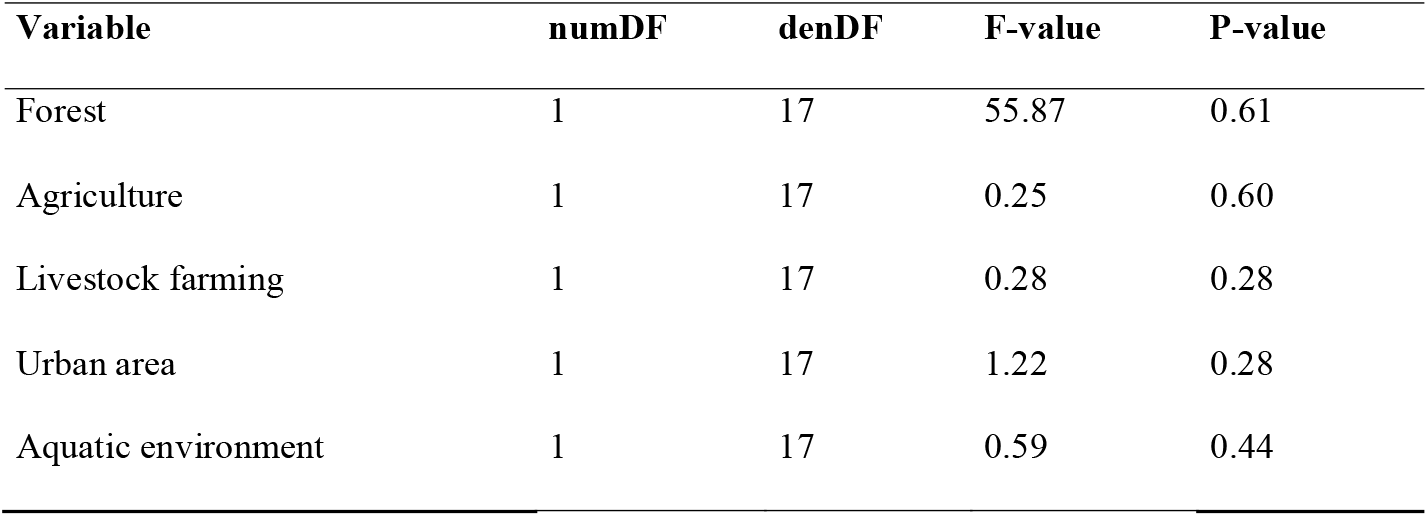
Result of the Generalized Linear Mixed-Effect Models (GLMM), with the function “glm” for amphibian richness and 250-m-buffer landscape variables in 23 water bodies (breeding sites) in southern Brazil, recorded from October 2018 to March 2019

**Fig. 4.**
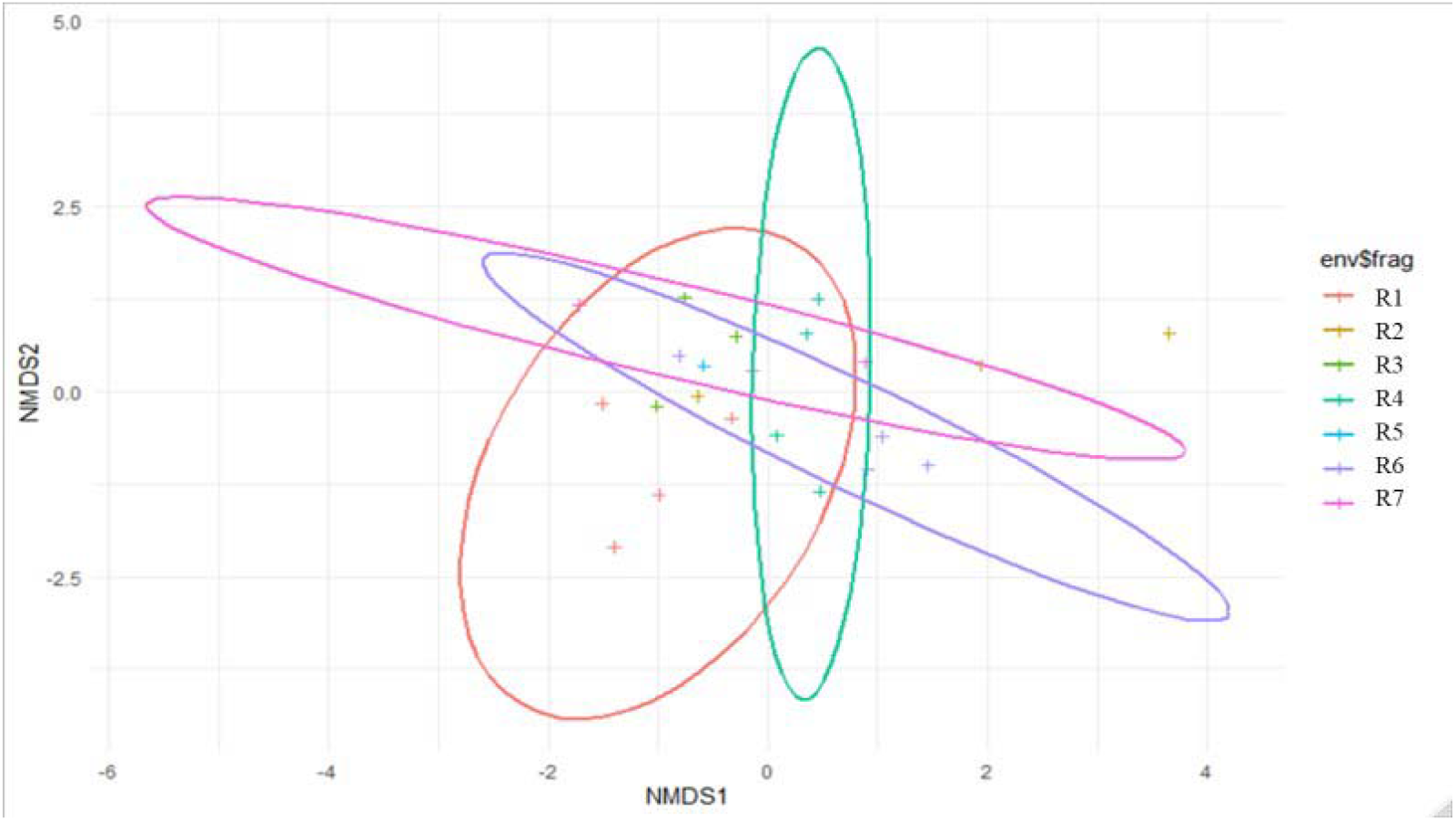
Non-metric multidimensional scaling (NMDS) based on Bray-Curtis distances and integrated environmental parameters depicting compositional differences among forest remnants. Only two dimensions are given to facilitate interpretation. The ellipses represent the formed groups, the symbols (+) represent the sampled breeding sites and the sampled forest remnants are represented as follows: + R1 = Parque Estadual de Vila Velha; + R2 = Parque Estadual Rio Guarani; + R3 = Reserva Privada Enele; + R4 = Parque Estadual das Araucárias; + R5 = Reserva Privada Quebra-queixo; + R6 = Parque Estadual Fritz Plaumann; + R7 = Parque Estadual do Turvo

Satellite image analysis showed that the dominant classes of land use were forests and livestock pasture (Table S2). Together, forests and livestock farming were pointed out by PERMANOVA as the main landscape component to explain the patterns of compositional dissimilarity (beta diversity) between remnants (forest, R^2^ = 0.10, F = 2.35, p = 0.01; livestock farming, R^2^ = 0.08, F = 1.93, p = 0.02; see Table 3).

**Table 3.** Results of PERMANOVA showing the contribution of landscape variables to the dissimilarity patterns in amphibian composition in the set of communities

| Variable | df | Sums Sqs | Mean Sqs | F-Model | R2 | P-value |
| --- | --- | --- | --- | --- | --- | --- |
| Forest | 1 | 0.91 | 0.91 | 2.35 | 0.09 | 0.01* |
| Agriculture | 1 | 0.59 | 0.59 | 1.52 | 0.06 | 0.39 |
| Livestock farming | 1 | 0.75 | 0.75 | 1.93 | 0.08 | 0.02* |
| Urban area | 1 | 0.38 | 0.38 | 0.98 | 0.04 | 0.22 |
| Aquatic environment | 1 | 0.28 | 0.28 | 0.74 | 0.03 | 0.34 |
| Residuals | 17 | 6.60 | 0.38 |  | 1.00 |  |
| Total | 22 | 9.54 |  |  |  |  |

## Discussion

We recorded 22 anuran species, which correspond approximately to two-thirds of the richness found in similar forest remnants in southern Brazil (Lucas and Fortes 2008; Iop et al. 2012; Bastiani and Lucas 2013). Hylidae represented over half of species, most of which are considered generalists and found in both forest environments and grasslands or forest edges (Lucas and Fortes 2008; Almeida-Gomes and Rocha 2014; Barbosa et al. 2014; Oliveira et al. 2017). The lack of any strong association between tadpole species richness and land use suggests that anurans are primally affected by habitat characteristics which are detected only on a fine-scale analysis. Breeding site configuration, such as pond descriptors, for example, would play a bigger role than land use over tadpole species diversity (Dalmolin et al. 2020). Although we have no detailed data to support this hypothesis, several studies have shown that microhabitat characteristics affect amphibian richness (D’Anunciação et al. 2013; Knauth et al. 2018; Figueiredo et al. 2019). Forest heterogeneity, leaf litter depth, canopy cover and presence of clearings are examples of local descriptors that are not detected at a landscape level but affect anuran species composition (Ferrante et al. 2017; Howell et al. 2019). Most local elements of the habitats support species persistence by providing fundamental resources as well as shelter (Erős et al. 2014; Landeiro et al. 2014; Datry et al. 2016; Collins and Fahrig 2017). This idea is reinforced by the fact that *Boana curupi* and *Crossodactylus schimidti* were registered only where local characteristics, such as streams with a rocky bottom, were present (Bastiani et al. 2012; Bastiani and Lucas 2013; Caldart et al. 2013).

One possible explanation for the similarities in species composition between forest remnants is the fact that they are surrounded by relatively well-preserved landscapes. Although a little speculative, we believe that that they could offer favorable conditions for the maintenance of local populations and homogenize species composition at a regional scale. As the remnants are surrounded mainly by other forest formations and pastures, they are little exposed to pollution and other anthropogenic disturbs, being able to support high species diversity (Becker et al. 2010; Ferreira et al. 2016; Preuss et al. 2020).

However, we do not minimize the relevance of landscape proprieties, such as the amount of available habitat, for the colonization and persistence of species (Faggioni et al. 2020). Seasonal displacements, such as those related to mating and oviposition, involve many risks, and landscape changes could negatively affect them, causing a strong impact on reproductive cycles (Becker et al. 2010). Thus, we strongly recommend new landscape-scale studies evaluating anuran diversity in a wider range.

Despite the predominance of forested habitats, we recorded only two forest species (*Boana curupi* and *Crossodactylus schmidti*). Species composition was dominated by habitat generalists (*Boana faber, Dendropsophus minutus, Lithobates catesbeianus, Physalaemus* cf. *carrizorum, P. cuvieri, Scinax fuscovarius*). The predominance of habitat generalists and widely distributed species has already been pointed out in other forest remnants that were studied in this portion of the Atlantic Forest (Lucas and Fortes 2008; Lucas and Marocco 2011; Bastiani and Lucas 2013). This could be a result of the deforestation of the Atlantic forest, in which species adapted to open conditions replace specialized species that are adapted to the forest (Haddad and Prado 2005). Homogenization of biota (Ferrante et al. 2017; Nowakowski et al. 2018) could explain the lack of relationship between species composition and landscape properties that was recorded.

We highlight the fact that, in the whole sampled area, forest species were limited to two species (*Boana curupi* and *Crossodactylus schmidti*). We have no information about the species composition in the studied forest remnants in past decades. Thus, we cannot affirm whether we are showing a recent or a well-established scenario about the regional anuran species composition. Forest-related species tend to be more prone to population decline due to their low ability to migrate across areas where the canopy cover is absent (Howell et al. 2019). The conversion of forest to pasture, which occurred decades ago, could have favored the local extinction of other forest-related anuran species. At the same time, these habitat changes favor the colonization by species such as *Boana faber* and *Dendropsophus minutus* (Aquino et al. 2004, 2010; Scott et al. 2004; Lavilla et al. 2010; Silvano et al. 2010). These species are often found in fragmented landscapes and expanding open areas (Preuss 2018; Figueiredo et al. 2019; Menin et al. 2019). We also recorded the exotic bullfrog (*Lithobates catesbeianus*) in four of the seven remnants. This species is widely distributed and well established in altered environments of Brazil’s South region (Both et al. 2011; Madalozzo et al. 2016). The process of dispersion and colonization of new localities is possibly expanding (Santos-Pereira and Rocha 2015) together with anthropogenic changes. Little is still known about the persistence of populations in altered environments, so this is an important subject to be investigated in future studies.

The presence of livestock pasture (farming) and forests were the landscape components that best explained the dissimilarity in species composition. The role of forests and pastures over amphibian species composition is relatively well understood (Haddad et al. 2013; Howell et al. 2019). At the same time, the conversion of forests into areas of pasture, agriculture or urbanization may limit or prevent the permanence of forest-associated species. In this sense, the composition of amphibian communities in remaining natural areas may change over time as a result of the colonization or permanence of generalist species (Ferrante et al. 2017). In this sense, we encourage future studies that deal with past and present species composition and the role of landscape changes and different surrounding matrices in past decades for current species assemblages.

## Compliance with ethical standards

### Ethics approval

All applicable international, national, and/or institutional guidelines for the care and use of animals were followed.

### Consent to participate

Not applicable.

### Consent for publication

Not applicable.

## Acknowledgments

We are thankful to all private owners that authorized the entry into their properties and the state environmental agencies that authorized the research in the conservation units.

## Availability of data and material

All datasets of this study are available through the corresponding author on reasonable request.

## Code availability

Not applicable.

## Authors’ contributions

All authors contributed to the study conception and design, material preparation, and writing of the manuscript. Data collection and species identification were performed by RCS, remote sensory analysis was performed by DB and data analysis was performed by DAD. All authors read and approved the final manuscript.

