## Supplementary files for "Association between land use and composition of amphibian species in temperate Brazilian forest remnants"

**Table S1** Description of tadpole sample points in southern Brazil, from October 2018 to March 2019. VV = Parque Estadual de Vila Velha; RG = Parque Estadual Rio Guarani; RE = Reserva Privada Enele; PA = Parque Estadual das Araucárias; QQ = Reserva Privada Quebra-Queixo; FP = Parque Estadual Fritz Plaumann; PT = Parque Estadual do Turvo

| Water body | Study area | Sampling site | State | Geographic coordinates | Sample point | Description of the place |
|---|---|---|---|---|---|---|
| 1 | 1 | VV | PR | 25°13'44.23"S 50° 2'8.87"O | Pond | Pond, with grassy and aquatic vegetation, in secondary forest and close to a road, inside protected area. |
| 2 | 1 | VV | PR | 25°14'43.39"S 50° 0'59.01"O | Stream | Stream, with grassland and shrub formation, inside the protected area. |
| 3 | 1 | VV | PR | 25°14'52.85"S 49°59'30.90"O | Stream | Stream, with grassland and shrub formation, inside protected area. |
| 4 | 1 | VV | PR | 25°15'33.85"S 50° 1'26.77"O | Pond | Pond, with landscape of exposed soil, grass and shrubs close to secondary vegetation, outside the protected area. |
| 5 | 1 | VV | PR | 25°15'29.01"S 50° 1'20.75"O | Stream | Stream, with landscape of exposed soil, grass and shrubs, close to secondary vegetation, outside the protected area. |

| | | | | | | |
|---|---|---|---|---|---|---|
| 6 | 2 | RG | PR | 25°26'25.73"S<br>53° 9'52.66"O | Pond | Artificial pond, with the presence of secondary forest, at forest border, inside the protected area. |
| 7 | 2 | RG | PR | 25°26'20.44"S<br>53° 9'10.23"O | Stream | Stream, with the presence of secondary forest, inside the protected area. |
| 8 | 2 | RG | PR | 25°26'19.53"S<br>53°10'11.56"O | Pond | Artificial pond, with the presence of pasture and agriculture in the surroundings, outside the protected area. |
| 9 | 2 | RG | PR | 25°26'16.72"S<br>53°10'0.94"O | Stream | Stream, with the presence of riparian forest with secondary vegetation, outside the protected area. |
| 10 | 3 | RE | SC | 26°22'3.39"S<br>52°50'28.57"O | Pond | Artificial pond, with the presence of pasture, grass and aquatic plants, inside the private reserve. |
| 11 | 3 | RE | SC | 26°21'57.65"S<br>52°50'1.40"O | Stream | Second-order stream, with the presence of secondary forest, inside the private reserve. |
| 12 | 3 | RE | SC | 26°22'2.64"S<br>52°49'47.29"O | Pond | Artificial pond, with the presence of pasture, close to urban area, outside the private reserve. |
| 13 | 3 | RE | SC | 26°22'0.88"S<br>52°49'40.87"O | Stream | Stream spring, with the presence of pasture, grasses and shrubs, close to urban area, outside the private reserve. |

| | | | | | | |
|---|---|---|---|---|---|---|
| 14 | 4 | PA | SC | 26°27'27.94"S 52°33'47.77"O | Pond | Pond, with the presence of aquatic vegetation, grass and secondary forest in the surroundings, inside the protected area. |
| 15 | 4 | PA | SC | 26°27'28.93"S 52°33'45.54"O | Stream | Stream, with the presence of secondary forest, inside the protected area. |
| 16 | 4 | PA | SC | 26°28'8.27"S 52°34'17.88"O | Stream | Stream, with the presence of secondary forest, inside the protected area. |
| 17 | 4 | PA | SC | 26°29'1.15"S 52°33'18.43"O | Pond | Pond, with the presence of aquatic vegetation, grasses and secondary forest in the surroundings, pasture and crops, outside the protected area. |
| 18 | 4 | PA | SC | 26°27'6.36"S 52°33'34.69"O | Stream | Stream, with the presence of shrubby riparian forest and crops, outside the protected area. |
| 19 | 5 | QQ | SC | 26°39'7.80"S 52°32'30.37"O | Pond | Artificial pond, with the presence of aquatic vegetation, grasses, pastures in secondary forest and lake, outside the private reserve. |
| 20 | 5 | QQ | SC | 26°38'59.58"S 52°32'20.11"O | Stream | Stream, in secondary forest, close to pasture and lake, inside the private reserve. |
| 21 | 6 | FP | SC | 27°17'29.56"S 52° 6'42.63"O | Stream | Stream, with the presence of secondary forest, inside the protected area. |

| | | | | | | |
|---|---|---|---|---|---|---|
| 22 | 6 | FP | SC | 27°17'21.45"S<br>52° 6'5.13"O | Pond | Artificial pond, with the presence of aquatic vegetation, grasses and secondary forest in the surroundings, pasture and crops, outside the protected area. |
| 23 | 6 | FP | SC | 27°17'37.40"S<br>52° 6'15.14"O | Stream | Stream spring, with the presence of grasses and shrubs, outside the protected area. |
| 24 | 6 | FP | SC | 27°17'30.69"S<br>52° 5'32.35"O | Stream | Stream, with the presence of grasses and secondary riparian forest, outside protected area. |
| 25 | 7 | TP | RS | 27°13'28.07"S<br>53°51'6.13"O | Pond | Pond with aquatic vegetation, surrounded secondary forest and close to road. inside protected area. |
| 26 | 7 | TP | RS | 27°13'58.52"S<br>53°51'17.21"O | Stream | Lotic water, on the edge of secondary forest, inside protected area. |
| 27 | 7 | TP | RS | 27°15'4.95"S<br>53°56'20.40"O | Pond | Semi-temporary artificial pond, surrounded by grassy vegetation. Outside protected area, in agriculture. |
| 28 | 7 | TP | RS | 27°14'30.90"S<br>53°50'24.61"O | Pond | Semi-temporary artificial pond, surrounded by grassy vegetation. Outside protected area, in agriculture. |

**Table S2** Average value of landscape components (km²) in the sampled sites in a 250-m-radius buffer, in southern Brazil, from October 2018 to March 2019

| **Category of land use** | **Buffer occupied area (km²)** | | | | | | |
|---|---|---|---|---|---|---|---|
| | Remnant 1 | Remnant 2 | Remnant 3 | Remnant 4 | Remnant 5 | Remnant 6 | Remnant 7 |
| Agriculture | 0.00 | 0.00 | 0.00 | 0.04 | 0.00 | 0.07 | 0.08 |
| Aquatic environment | 0.00 | 0.00 | 0.00 | 0.00 | 0.05 | 0.00 | 0.00 |
| Forest | 0.09 | 0.10 | 0.08 | 0.13 | 0.00 | 0.13 | 0.07 |
| Livestock farming | 0.00 | 0.10 | 0.10 | 0.03 | 0.15 | 0.00 | 0.05 |
| Urban area | 0.01 | 0.00 | 0.01 | 0.00 | 0.00 | 0.00 | 0.00 |

**Table S3** Summary Results of Principal Components Analysis

| Axis | Eigenvalue variance | Variance percent | Percent cumulative |
|---|---|---|---|
| Axis.1 | 1.72 | **34.56** | 34.56 |
| Axis.2 | 1.37 | **27.52** | **62.08** |
| Axis.3 | 1.04 | 20.97 | 83.06 |
| Axis.4 | 0.66 | 13.28 | 96.35 |
| Axis.5 | 0.18 | 3.64 | 100 |

**Table S4** Correlations between communities and landscape variables with the axes of PCA

| **Descriptors** | **Axis 1** | **Axis 2** | **Axis 3** | **Axis 4** | **Axis 5** |
|---|---|---|---|---|---|
| Forest | -0.84 | 0.43 | -0.15 | 0.06 | -0.27 |
| Agriculture | 0.09 | -0.94 | 0.12 | 0.15 | -0.23 |
| Livestock farming | 0.81 | 0.31 | 0.01 | -0.42 | -0.21 |
| Urban area | 0.24 | 0.38 | 0.78 | 0.41 | -0.04 |
| Aquatic environment | 0.53 | 0.18 | -0.62 | 0.53 | -0.04 |
| Community 1 | -1.14 | 0.92 | 5.42 | 0.64 | -0.19 |
| Community 2 | 0.13 | -0.33 | 5.27 | -0.11 | 1.46 |
| Community 3 | 0.04 | 0.23 | 8.15 | 0.50 | 1.01 |
| Community 4 | -0.72 | 0.17 | -2.00 | -0.01 | 0.57 |
| Community 5 | 0.28 | 0.53 | -1.75 | -0.81 | -0.11 |
| Community 6 | -1.43 | 0.58 | -3.63 | 0.07 | -0.14 |
| Community 7 | 2.01 | 0.48 | 1.33 | -1.70 | -0.09 |

| | | | | | |
|---|---|---|---|---|---|
| Community 8 | 1.49 | 0.50 | -4.32 | -1.43 | -0.09 |
| Community 9 | -1.09 | 0.57 | -3.26 | -0.10 | -0.14 |
| Community 10 | 1.82 | 1.86 | 3.48 | 1.30 | -0.29 |
| Community 11 | -1.25 | 0.24 | -2.88 | 0.11 | -0.15 |
| Community 12 | -1.43 | 0.58 | -3.63 | 0.07 | -0.14 |
| Community 13 | 0.64 | 0.18 | -8.11 | -0.86 | -0.12 |
| Community 14 | -0.26 | -1.64 | 1.24 | 0.34 | -0.19 |
| Community 15 | 3.24 | 1.01 | -2.93 | 1.98 | -0.07 |
| Community 16 | -1.43 | 0.58 | -3.63 | 0.07 | -0.14 |
| Community 17 | 0.35 | -2.85 | 3.87 | 0.49 | -0.21 |
| Community 18 | -0.62 | -0.95 | -2.54 | 0.26 | -0.17 |
| Community 19 | -1.43 | 0.58 | -3.63 | 0.07 | -0.14 |
| Community 20 | -1.43 | 0.58 | -3.63 | 0.07 | -0.14 |
| Community 21 | 0.98 | 0.51 | -9.97 | -1.17 | -0.10 |
| Community 22 | 0.69 | -2.18 | 3.13 | 0.05 | -0.19 |
| Community 23 | 0.59 | -2.22 | 3.12 | 0.13 | -0.11 |
